# The zebrafish *goosepimples/myosin Vb* mutant exhibits cellular attributes of human microvillus inclusion disease

**DOI:** 10.1101/030536

**Authors:** Jaydeep Sidhaye, Clyde Pinto, Shweta Dharap, Tressa Jacob, Shobha Bhargava, Mahendra Sonawane

**Author notes:** Current Address: Max-Planck-Institute of Molecular Cell Biology and Genetics, Dresden, Germany (Jaydeep Sidhaye). These authors contributed equally (Jaydeep Sidhaye, Clyde Pinto). Bullet points (Key findings): - *myosin Vb* is expressed in the zebrafish intestine. - *goosepimples/myosin Vb* function is essential for the morphogenesis, cell shape modulation, secretion of glycoproteins and prevention of the formation of microvillus inclusions in the zebrafish intestine. - The function of *myosin Vb* in the intestine is conserved between fish and humans. - The *goosepimples* mutant is a good animal model for microvillus inclusion disease in humans. Grant Sponsor: Wellcome Trust-DBT India alliance (MS; 500129-Z-09-Z), Department of Science and Technology, India (MS; SR/S2/RJN-06/2006)’ TIFR-DAE (MS; 12P-121).

## Abstract

**Background:** Microvillus inclusion disease (MVID) is a life threatening enteropathy characterised by perpetual diarrhoea. Recent analysis has revealed that enterocytes in MVID patients exhibit reduction of microvilli, presence of microvillus inclusion bodies and intestinal villus atrophy. Genetic linkage analysis has identified mutations in *myosin Vb* gene as the main cause of MVID. An animal model that develops ex-utero and is tractable genetically as well as by microscopy would be highly useful to study the cellular basis of the formation of inclusion bodies.

**Results:** Here we report that the intestine of the zebrafish *goosepimples (gsp)/myosin Vb (myoVb)* mutant show severe reduction in the intestinal folds-structures similar to mammalian villi. The loss of folds is further correlated with changes in the shape of enterocytes. In a striking similarity with MVID patients, zebrafish *gsp/myoVb* mutant larvae exhibit microvillus atrophy, microvillus inclusions and accumulation of secretory material in enterocytes.

**Conclusion:** We propose that the zebrafish *gsp/myoVb* mutant is a valuable model to study the pathophysiology of MVID. Owing to the advantages of zebrafish in screening libraries of small molecules, the *gsp* mutant will be an appropriate tool to identify compounds that would alleviate the formation of microvillus inclusions and have therapeutic value.

## Introduction

Microvillus inclusion disease (MVID) is a dreadful enteropathy inherited in an autosomal recessive manner and reported in infants often born of consanguineous parents. The main symptom of MVID is intractable diarrhoea. While the exact biological reason for the watery diarrhoea is not clear, the classic features of MVID at the cellular and tissue level have been well described. These include villus atrophy, reduction or shortening of microvilli and the presence of inclusion bodies in patient enterocytes (Golachowska et al., 2010). In addition to inclusion bodies, altered distribution of apical markers such as sucrose isomaltase (Ameen and Salas, 2000) and accumulation of Periodic acid-Schiff staining (PAS) positive material, presumably secretory granules (Phillips et al., 2004), have also been reported in the enterocytes of MVID patients. The survival of MVID patients depends mainly on long-term parenteral nutrition. However, long-term parenteral nutrition may involve secondary complications such as bacterial infections, cholestatic liver disease and vascular as well as renal complications. The only successful MVID remedy so far has been bowel transplant (Oliva et al., 1994; Bunn et al., 2000; Ruemmele et al., 2004). This necessitates further investigation of the cellular and physiological basis of the disease and possible remedies.

Homozygosity mapping using single nucleotide polymorphisms (SNPs) in Turkish kindred has unravelled that mutations in the *myosin Vb* gene are a major cause of MVID (Muller et al., 2008). By now, multiple studies have shown the association between mutations in *myosin Vb* and MVID (Muller et al., 2008; Ruemmele et al., 2010; Szperl et al., 2011)). A recently prepared international patient registry reports 41 unique *myosin Vb* mutations thought to be associated with MVID (van der Velde et al., 2013). Myosin Vb is an actin based molecular motor. It interacts with Rab11, Rab8 and Rab10 and has been shown to be involved in transport of recycling endosomes and plasma membrane biogenesis (Lapierre et al., 2001; Roland et al., 2007; Liu et al., 2013). Consistently, it has been shown that loss of *myosin Vb* function results in the loss of Rab11-FIP5 positive recycling endosomes in the enterocytes of MVID patients (Szperl et al., 2011). Furthermore, the apical transporter protein CD36 and basolateral transporters such as the Na^+^/K^+^ ATPase and transferrin receptor are mislocalised in MVID patients, suggesting that some aspects of enterocyte polarisation are compromised (Muller et al., 2008; Thoeni et al., 2014).

All studies involving the physiological and cellular characterisation of this disease depend on clinical explants obtained through biopsies of patients. So far, the knockdown of *myosin-Vb* using si-RNA in the Caco2 cell line has been used as a model for MVID (Ruemmele et al., 2010; Thoeni et al., 2014). Very recently, a Myo5b knockout mouse model exhibiting attributes of MVID has been published (Carton-Garcia et al., 2015). Other animal models for this disease include *Rab8, Rab11a* and *Cdc42* knockout mice, which also exhibit microvillus inclusions in the enterocytes (Sato et al., 2007; Sakamori et al., 2012; Sobajima et al., 2014). A vertebrate model that is tractable by light and fluorescence microscopy would be an asset in understanding the cell biology of MVID and unravelling the role of Myosin Vb mediated trafficking in the intestinal epithelium.

Over the last 20 years, zebrafish has emerged as a powerful model to study human diseases and disorders (Ingham, 2009). Zebrafish disease models are useful to understand the pathophysiology of a disease and to screen libraries of small molecules that may have therapeutic value (Peterson and Fishman, 2011). The zebrafish larval intestine is easily visible under a stereomicroscope through the transparent skin. Even at early larval stages it shows gross morphological similarities with the human intestine. The similarities at the cellular, molecular and functional level are quite remarkable (Wallace et al., 2005). The intestine of a zebrafish exhibit folds, which are similar to mammalian villi, while its enterocytes exhibit apical microvilli just like mammalian enterocytes. Thus, the zebrafish intestine provides an excellent model to investigate intestinal development and disease.

Here we report that the intestines of zebrafish *gsp/myoVb* mutant larvae exhibit the classic features of MVID. The mutant enterocytes contain microvillus inclusion bodies very similar to MVID patients. There is also a reduction in the length of microvilli in the mutants and the shape of the mutant enterocytes is altered, possibly affecting the formation of intestinal folds. Furthermore, we show that the mutants have defects in lipid absorption. Our analyses presented here indicate that the function of Myosin Vb is conserved between fish and humans and that the zebrafish *gsp* mutant is an excellent animal model to study the cellular basis of MVID.

## Results

### *gsp/myoVb* is expressed in the developing zebrafish gut

We have recently shown that the *gsp* locus, which is essential for the maintenance of epidermal architecture, encodes for the actin based molecular motor (Sonal et al., 2014). While the morphological phenotype in the epidermis subsides after 3 days post fertilisation (dpf), the mutants typically die at the beginning of metamorphosis due to unknown reasons. Keeping in mind the association of *myoVb* with MVID, we asked if the reason for death could be linked to an intestinal defect. We checked the expression of *myoVb* in the developing zebrafish gut by performing in situ hybridisation. Our analysis revealed that *myoVb* is predominantly expressed in the anterior part of the intestine or intestinal bulb between 3 and 6 dpf. While considerable *myoVb* expression was also seen in the proximal part of the midgut, we did not observe its expression in the distal midgut and posterior gut using in situ hybridisation (Fig. 1A-O). RT-PCR analysis on isolated zebrafish intestines from 4-day-old larvae revealed significant expression of *myoVb* as well as two other *myosin V* paralogues in the zebrafish intestine (Fig. 1P).

**Figure 1:**
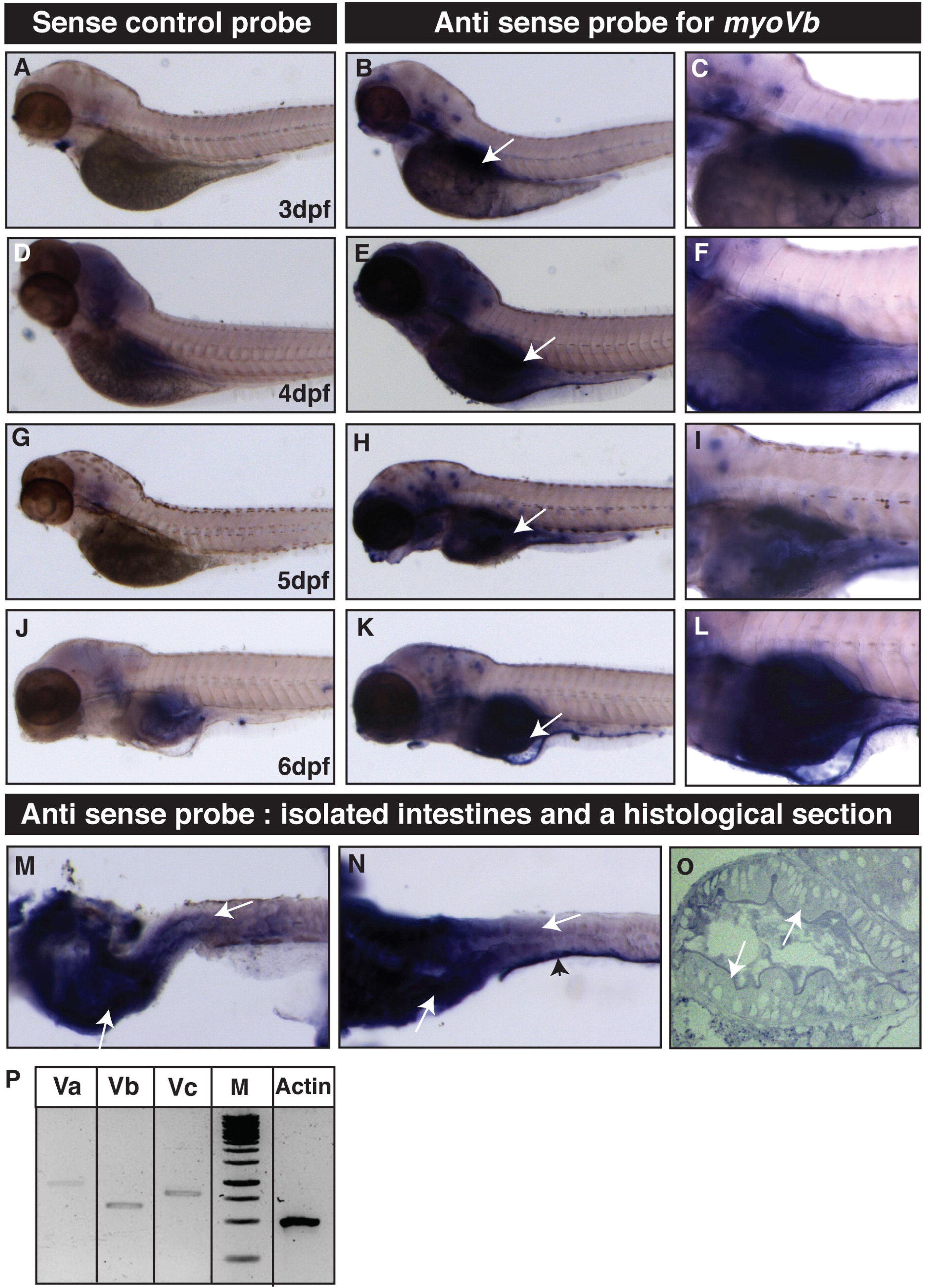
Temporal expression analysis reveals that *gsp/myoVb* is expressed in the intestine of zebrafish larvae. In situ hybridisation analyses using sense control probe (A, D, G, J) and antisense probe against *myosin Vb* at lower magnification (B, E, H, K) as well as higher magnification (C, F, I, L) reveal that *myosin Vb* is expressed in the intestine during day 3 to 6 of zebrafish larval development. Note that the isolated gut tissues (M, N) and the histological section (O) show clear presence of *myosin Vb* transcripts in the anterior gut and proximal midgut. RT-PCR analysis (P) shows that all three *myosin V* paralogues are expressed in the intestine at 4dpf. Abbreviation: M-1kb ladder marker. The white arrows point towards the expression of *myoVb.* The arrowhead points to the staining in the skin piece attached to the dissected gut.

We conclude that three *myosin V* paralogues are expressed in the developing zebrafish intestine. Furthermore, the expression of *myoVb* is observed in the intestinal bulb as well as in the proximal part of the midgut.

### The loss of *gsp/myoVb* function leads to a severe reduction in intestinal folds and concomitant changes in the shape of enterocytes

Since *myoVb* is expressed in the gut epithelium, we asked whether there is a morphological phenotype in the intestine of the *gsp/myoVb* mutant. Careful observation under a stereomicroscope revealed altered morphology of the intestine in the *gsp/myoVb* mutants as compared to wild type larvae. As reported earlier (Ng et al., 2005; Wallace et al., 2005), in wild type larvae, intestinal fold morphogenesis started at 4 dpf and subsequently the folds became more prominent (Fig. 2A,C,E,G,I,K). However, the mutant intestine lacked intestinal folds and appeared smooth-walled, even at 6 dpf (Fig. 2B,D,F,H,J,L). The observations were further substantiated by histological analysis performed on 6-day-old larvae (Fig 2M). In addition, the histological sections revealed the presence of goblet cells in the distal part of the midgut, indicating proper gut differentiation in the mutant larvae. As compared to wild type larvae, there was no major effect on peristalsis, albeit they appeared weaker in a few *gsp/myoVb* mutant larvae (data not shown).

**Figure 2:**
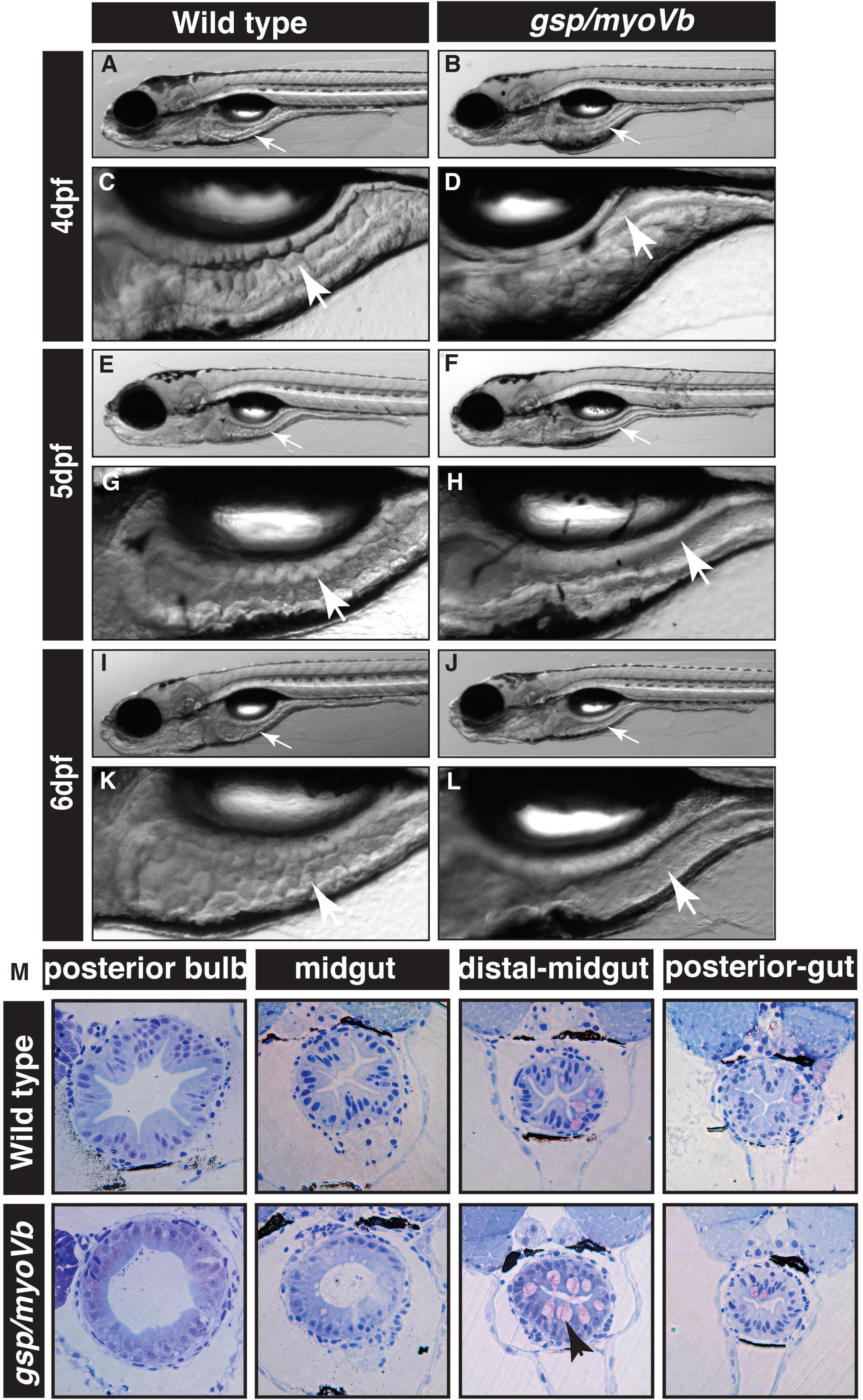
The zebrafish *gsp/myoVb* mutant exhibits smooth intestine phenotype during 4-to-6 days post-fertilisation. The bright field images of developing zebrafish wild type larvae (A, E, I) and their intestines (C, G, K) exhibit rough intestinal walls indicating the presence of tissue folds. The *gsp/myoVb* mutant larvae (B, F, J) and their intestines (D, H, L) show smooth intestinal walls suggesting absence of intestinal folds. Arrows point to the intestine in all the images. Histological sections (M) of different parts of the gut of wild type sibling and *gsp/myoVb* mutant. Total 4 wild type siblings and 4 mutants were analysed by histology.

To further confirm the effect on the formation of intestinal folds and to understand the possible cause of the phenotype, we asked whether the cell shape and cell polarisation is perturbed in the *gsp* mutant enterocytes. To analyse the shapes of enterocytes, we sectioned the gut from 6-day old wild type and *gsp/myoVb* mutant larvae in the background of *Tg*(*cldnB:lynEGFP*) transgenic line. In this transgenic line, *claudin B* promoter drives expression of lyn tagged EGFP in several epithelial tissues including the gut (Haas and Gilmour, 2006). We performed immunohistology on the wild type and mutant gut using anti GFP antibody. Corroborating the histology analysis, we observed a clear reduction in fold formation in the intestine of the *gsp* mutant larvae as compared to wild type (Fig. 3B-I). Furthermore, in wild type larvae, the crests of folds are formed by wedge shaped enterocytes with expanded/broad apical surfaces facing the lumen and narrow basolateral sides facing the extracellular matrix. In the *gsp/myoVb* mutants, the enterocytes appeared more columnar as well as taller as compared to the wild type (Fig. 3J,K). We further measured the central height, apical width and basal width of the intestinal cells from the histological sections of anterior, mid and posterior gut. This quantification confirmed that while the wild type enterocytes (7 larvae, n=88) are broader apically and narrower on the basal side, the *gsp/myoVb* mutant enterocytes (9 larvae, n=93) are apically narrower and basally broader (Fig. 3L; p<0.001). Additionally, the mutant enterocytes are taller than the enterocytes in wild type larvae (Fig. 3L; p<0.001).

**Figure 3:**
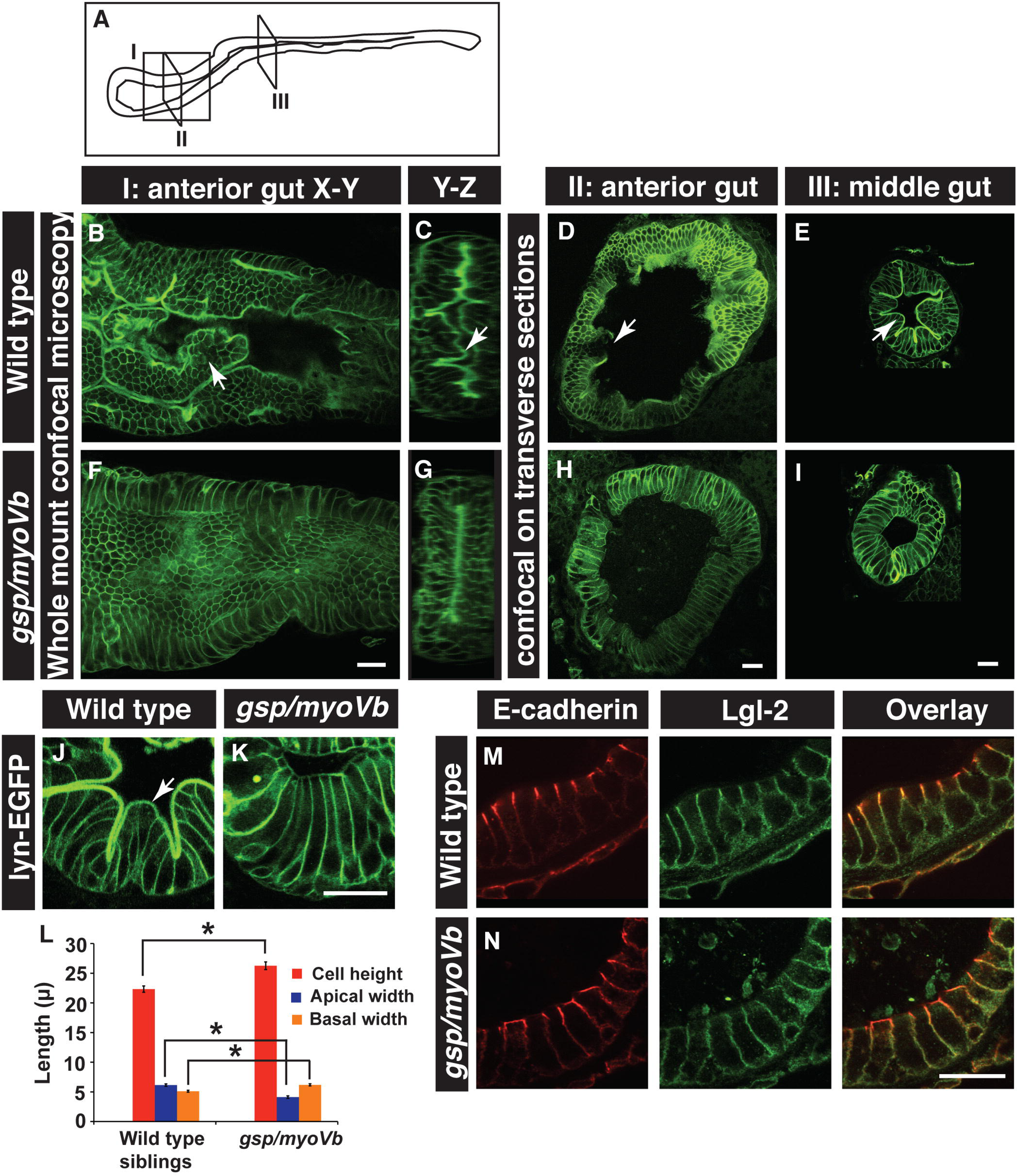
The loss of intestinal folds in *gsp/myoVb* mutants is accompanied by changes in cell shapes. A schematic showing various regions of the zebrafish larval gut (A). The boxed regions indicate the parts of the intestine where whole mounts were imaged (I) or intestines were sectioned (II, III). Confocal images of the whole mount (B,F), orthogonal sections (C,G), immuno-histological sections (D,E,H,IJ,K) of wild type siblings (B,C,D,EJ) and *gsp/myoVb* mutant (F,G,H,I,K) intestine stained for lyn-EGFP at 6dpf. Note the absence of intestinal folds in the *gsp/myoVb* mutants. The whole mount analysis was done on 4 siblings and 3 mutants whereas the immune-histology analysis was done on 9 siblings and 7 mutants. Bar graph (L) showing measurements of the height, apical width and basal width of enterocytes in the wild type and *gsp/myoVb* mutant larvae at 6 dpf. The enterocytes are taller and have a narrower apical domain in the *gsp/myoVb* mutants as compared to wild type. Immuno-histological sections of zebrafish intestine in the wild type (M) and *gsp/myoVb* mutant larvae (N) stained using anti E-cadherin and anti-Lgl2 antibodies. The overall polarity is maintained in the *gsp/myoVb* enterocytes.Arrows in B-E, J point to the intestinal folds. The square brackets show the comparison and asterisks indicates the statistically significant difference at p<0.05 by Student’s t-test. The error bars in L represent SEM. Scale bars are equivalent to 20μm.

Previous studies on MVID patient biopsies and the recent analysis of the *Myo5b* knockout mouse have indicated that certain transporter proteins, such as CD36, Na^+^/K^+^ATPase and transferrin receptor, are mislocalised in the absence of *myoVb* function, indicative of a partial disruption of apico-basal polarity in mature enterocytes (Muller et al., 2008; Thoeni et al., 2014; Carton-Garcia et al., 2015). We asked whether cell polarity is disrupted along with cell shapes in the enterocytes of *gsp* mutant larvae. Since the loss of or perturbations in cell polarity affect adherens junction formation, we analysed the localisation of E-cadherin — a transmembrane core component of the adherens junctions — in enterocytes. We used Lethal giant larvae 2 (Lgl2) as a marker for the basolateral domain. In siblings, E-cadherin localises laterally and is enriched on the apical side of the lateral surface, whereas Lgl2 localises to the basolateral domain. We did not observe any obvious defect in the localisation of E-cadherin or Lgl2 in the *gsp/myoVb* mutant intestine (Fig. 3M,N).

To conclude, we show that in the absence of *gsp/myoVb* function, intestinal folds do not form properly. Since cell shape change is one of the driving forces underlying morphogenesis, we propose that the effect of the loss of *gsp/myoVb* function on fold formation is manifested through the change in the enterocyte morphology. We further conclude that in the absence of MyoVb function, the gross apico-basal polarity of enterocytes is not perturbed in zebrafish.

### The *gsp/myoVb* mutant gut exhibits classic cellular features of MVID

In humans, mutations in the *myosin Vb* gene have been linked to MVID (Muller et al., 2008; Ruemmele et al., 2010). Analysis of intestinal cells of MVID patients has shown the presence of inclusion bodies with microvillar components trapped within (Ameen and Salas, 2000; Reinshagen et al., 2002; van der Velde et al., 2013; Thoeni et al., 2014). Enterocytes in Rab8a, Myo5b knockout mice and Myo5b deficient Caco2 cells exhibit the presence of actin rings marked by phalloidin, a classic feature of MVID (Sato et al., 2007; Ruemmele et al., 2010; Carton-Garcia et al., 2015). To check if such inclusion bodies are present in the enterocytes of *gsp/myoVb* mutant larvae, intestines of 6 dpf larvae were stained with phalloidin. Both sibling and mutant intestines showed bright apical actin staining corresponding to microvilli or apical brush border of the enterocytes (Fig. 4A,B). Strikingly, a few *gsp/myoVb* enterocytes showed the presence of large phalloidin labelled subapical vesicular bodies (Fig. 4B). These inclusion bodies have a diameter of about 3–5 μm. Similar inclusion bodies were also seen when intestinal sections were stained with phalloidin (Fig. 4C-D2). The inclusion bodies consisted of F-actin and are membrane bound. They are mostly associated with the apical membrane or are contiguous with the apical domain, suggesting their origin through invagination of the apical membrane (Fig. 4D2). We attempted to ascertain if there was a preponderance of these phalloidin rings in any particular region/s of the intestine. On visual inspection of phalloidin stained samples, rings were rarely observed in the posterior gut and the distal midgut (the region having a high density of goblet cells) so we restricted our analysis to the anterior gut and proximal midgut. Paired analysis revealed that phalloidin rings are more prevalent in the proximal midgut than in the anterior gut (Fig. 4E-G).

**Figure 4:**
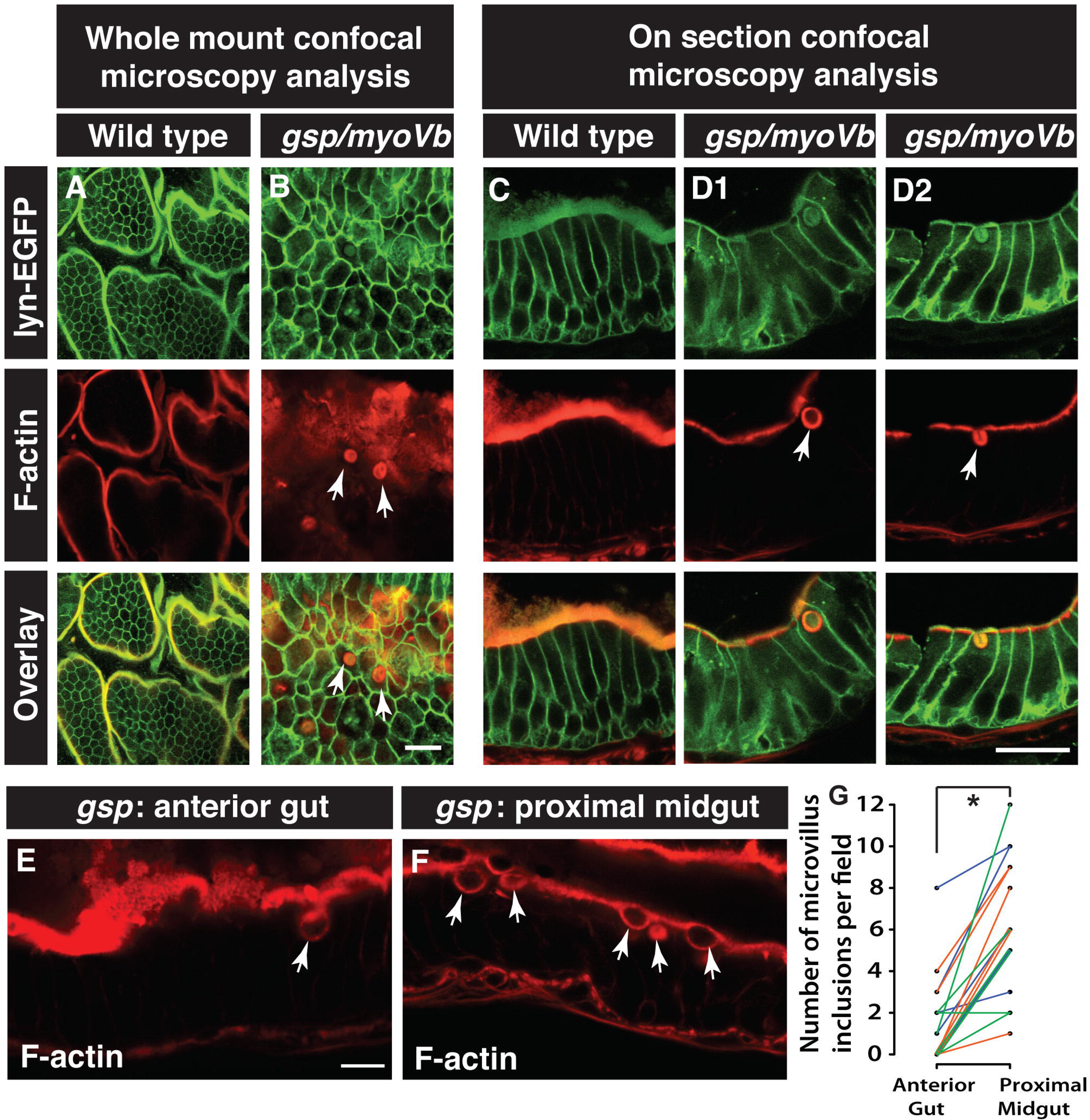
Enterocytes in *gsp/myoVb* mutants show presence of microvillus inclusions. Confocal microscopy analysis of whole mount (A,B) and cryosections (C,D1,D2) of the anterior gut of wild type sibling (A, C) and *gsp/myoVb* mutant (B,D1,D2) larvae stained for lyn-EGFP and F-actin (phalloidin) at 6dpf. In all, 3 mutants and 4 siblings were examined in whole mounts whereas 7 mutants and 7 siblings were analysed by cryosectioning. Confocal sections of anterior gut (E) and proximal midgut (F) from the same *gsp/myoVb* mutant stained for phalloidin. Total 17 mutant animals from three different sets were analysed this way. The graph (G) shows the comparison between the number of inclusions in the anterior versus proximal midgut in mutants. Each line indicates one animal and different colours indicate values from different experimental sets. The arrows indicate phalloidin or lyn-EGFP labelled inclusion bodies in enterocytes. The asterisk represents statistically significant difference (p<0.0001) by paired t-test. Scale bars: in B and D2 = 20 μm and in E=10 μm.

To confirm the presence of microvillus inclusions and to examine microvillus length, we performed TEM analysis on the intestines of wild type and mutant larvae at 6 dpf. In *gsp/myoVb* mutant enterocytes, there was a significant reduction in the length of microvilli as compared to the wild type (Fig. 5A-D, H). We also noticed that the extent of the reduction varied in different mutant animals (Fig. 5A-D, H). In addition, we observed microvillus inclusions in the mutant enterocytes at various subcellular locations such as at the apical side, at the lateral membrane domains and near the nucleus (Fig. 5B, E-G).

**Figure 5:**
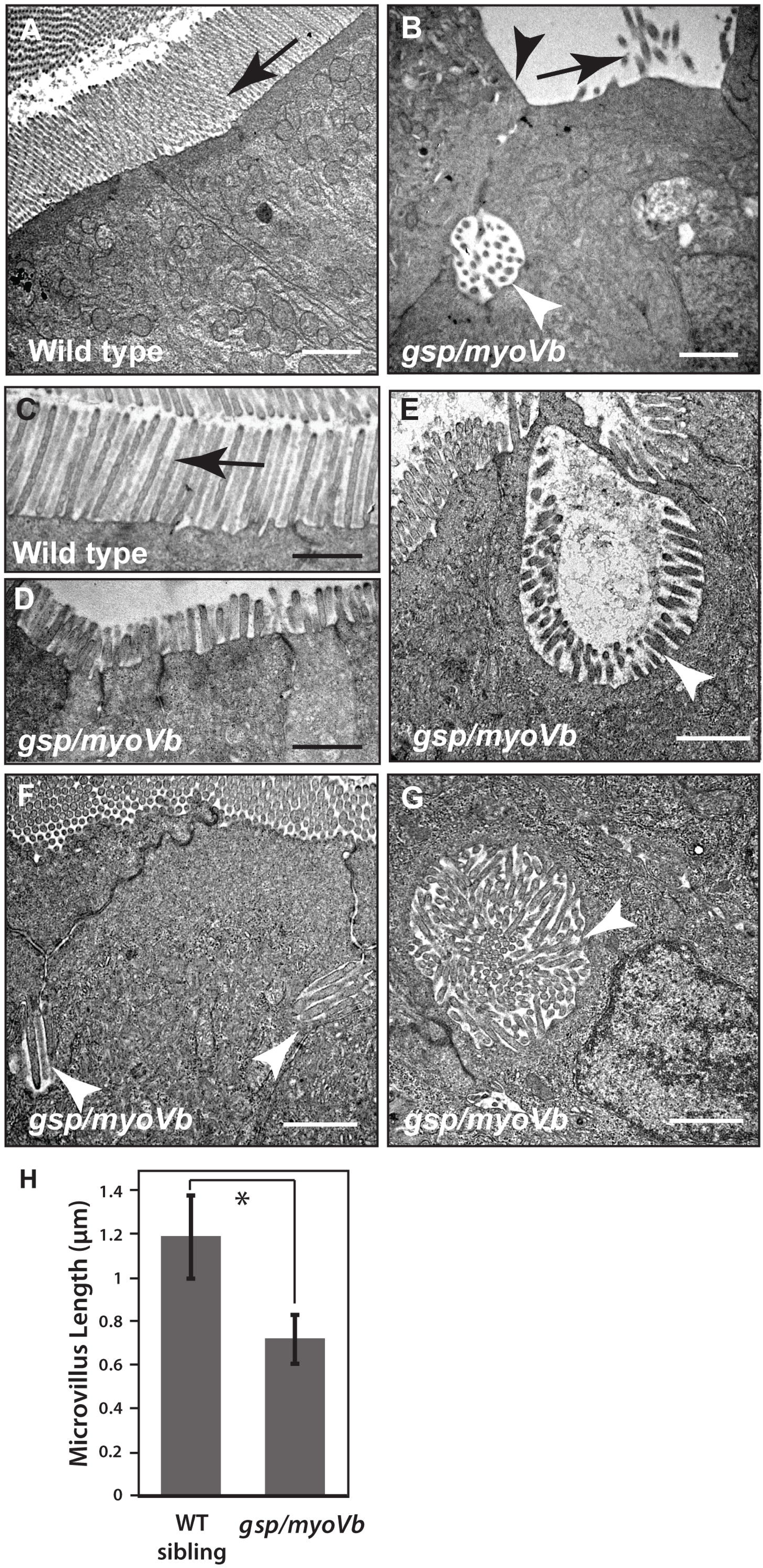
Shorter microvilli and microvillus inclusions, the main attributes of MVID, are present in enterocytes of *gsp/myoVb* mutants. Electron micrograph of thin sections, passing through the enterocytes of 6dpf wild type (A,C) and *gsp/myoVb* mutant larvae (B, D-G). Note the absence of microvilli (B), the reduction in microvillus length (D) and presence of microvillus inclusions (B, E-G) in *gsp/myoVb* enterocytes as compared to wild type enterocytes (A). The bar graph (H) shows the difference in the microvillar length in the midgut of wild type siblings and *gsp* mutants. For the EM analysis, we used 4 siblings and 5 mutant animals. The length measurements were done on 137 microvilli from 49 enterocytes of 4 siblings and on 95 microvilli from 33 enterocytes of 3 mutants. The white arrowheads indicate microvillus inclusions whereas the black arrows point to microvilli. A black arrowhead in (B) points to the loss of microvilli. The square bracket shows the comparison and asterisk indicates that the difference is statistically significant by an unpaired t-test with unequal variance at p<0.0001. The error bars in H represent the SD. Scale bars correspond to 1μm.

Another prominent feature of MVID is the accumulation of PAS positive material — presumably secretory granules — in the enterocytes (Phillips et al., 2004). PAS or Periodic acid-Schiff staining typically detects carbohydrates and glycoproteins. We asked whether *gsp/myoVb* mutants exhibit any such accumulation of secretory granules in the enterocytes. Instead of using PAS, we used fluorescently labelled Wheat Germ Agglutinin (WGA), which also binds to carbohydrates and glycoproteins. Our analysis performed on sections revealed large accumulation of glycoproteins in the apical domain of enterocytes confirming MVID-like cellular phenotypes in the *gsp/myosin Vb* mutant larvae (Fig. 6A,B).

**Figure 6:**
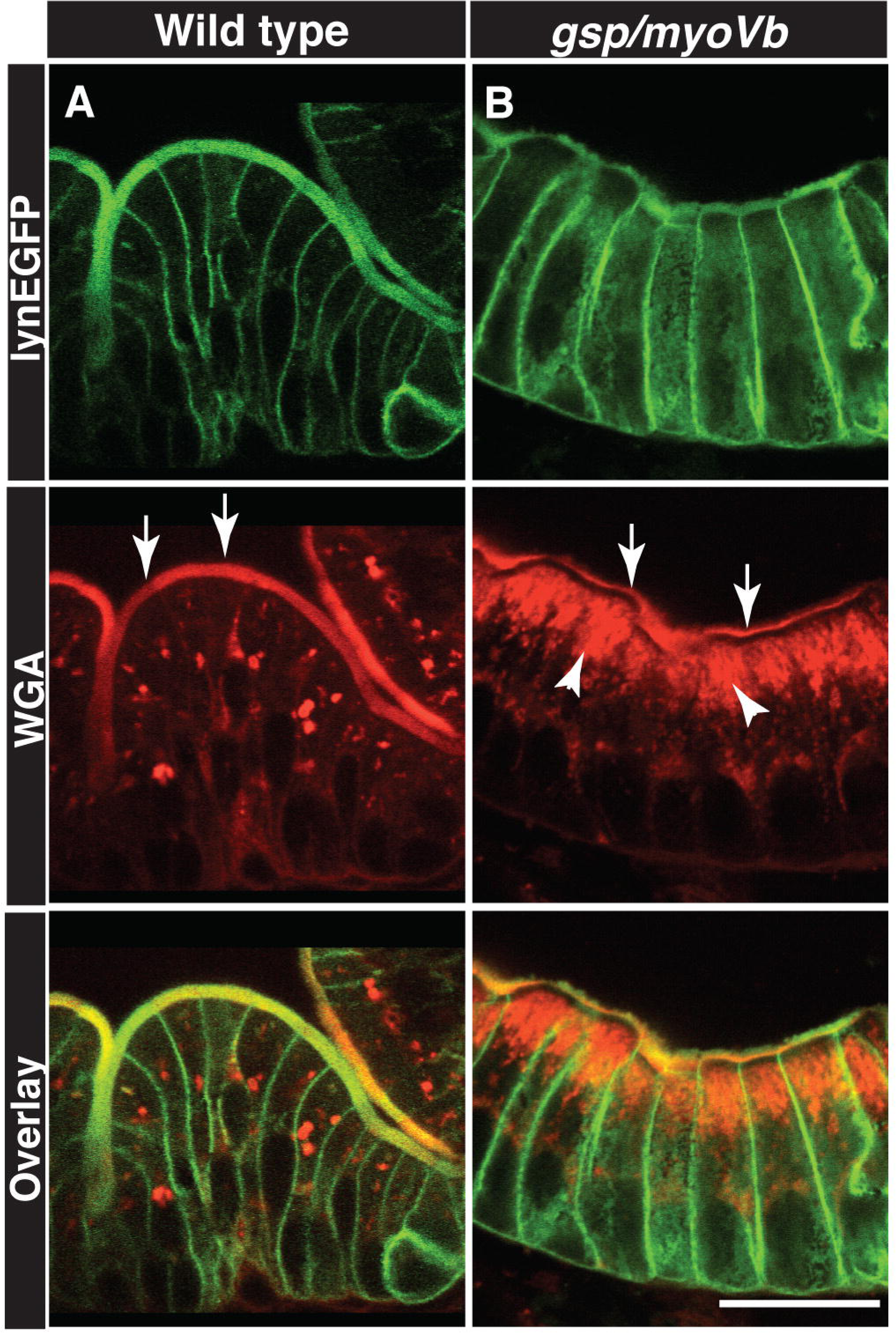
Secretory material accumulates in the enterocytes of *gsp/myoVb* mutant larvae. Immunohistology on 6dpf wild type siblings (A) and *gsp/myoVb* mutant (B) intestine using anti GFP antibody and Alexa fluor 594 conjugated WGA. The white arrows point to the WGA labelling of apical domain of the wild type and mutant enterocytes whereas the arrowheads point to the intracellular staining. Note the considerable decrease in apical staining and concomitant increase in intracellular staining in *gsp/myoVb* mutant. For this analysis, we checked 3 siblings and 4 mutants by histology. The scale bar is equivalent to 20 μm.

To conclude, histological as well electron microscopy analyses show that *gsp/myoVb* mutant enterocytes exhibit important attributes of MVID, such as reduction in microvillus length, presence of microvillus inclusions and accumulation of secretory granules.

### Lipid absorption is diminished in the *gsp/myoVb* mutant gut

Next, we asked if the cellular phenotypes in the *gsp/myoVb* mutant intestine also affect the absorptive function of the intestine. To test this directly, we incubated 6-day-old mutant and wild type larvae with either solid (boiled) or liquid (raw) egg-yolk, emulsified in fish water. To assess the uptake of food, an edible colouring agent was added in the yolk emulsion. We observed varied uptake of egg yolk by both wild type siblings as well as by the mutant larvae (Fig. 7A-D, data not shown). In the next set of experiments, which did not involve the colouring agent, we assayed the fed and unfed larvae for the presence of lipid droplets in the enterocytes (Walters et al., 2012). Electron microscopy and histological analysis on wild type siblings fed with either solid or liquid yolk revealed striking accumulation of lipid droplets in the enterocytes. In contrast, the *gsp/myoVb* mutant enterocytes exhibited a mild to drastic reduction in lipid accumulation (Fig. 7E-J; data not shown). Despite the variations observed in the extent of feeding, there was a consistent reduction in the lipid accumulation in the enterocytes of 9 mutant larvae, observed by histology (Data not shown). In the second set, we sorted the animals depending upon the consumption of coloured liquid yolk. Based on the colour intensity in the intestine, larvae were classified as ‘well-fed’ and ‘moderately-fed’ (Fig 7A-D). Of these, three well-fed wild type siblings and four well-fed mutants were further examined by histology. This analysis clearly revealed that well-fed mutant enterocytes exhibited reduced amount of lipid droplets as compared to the well-fed siblings (Fig. 7K,L)

**Figure 7:**
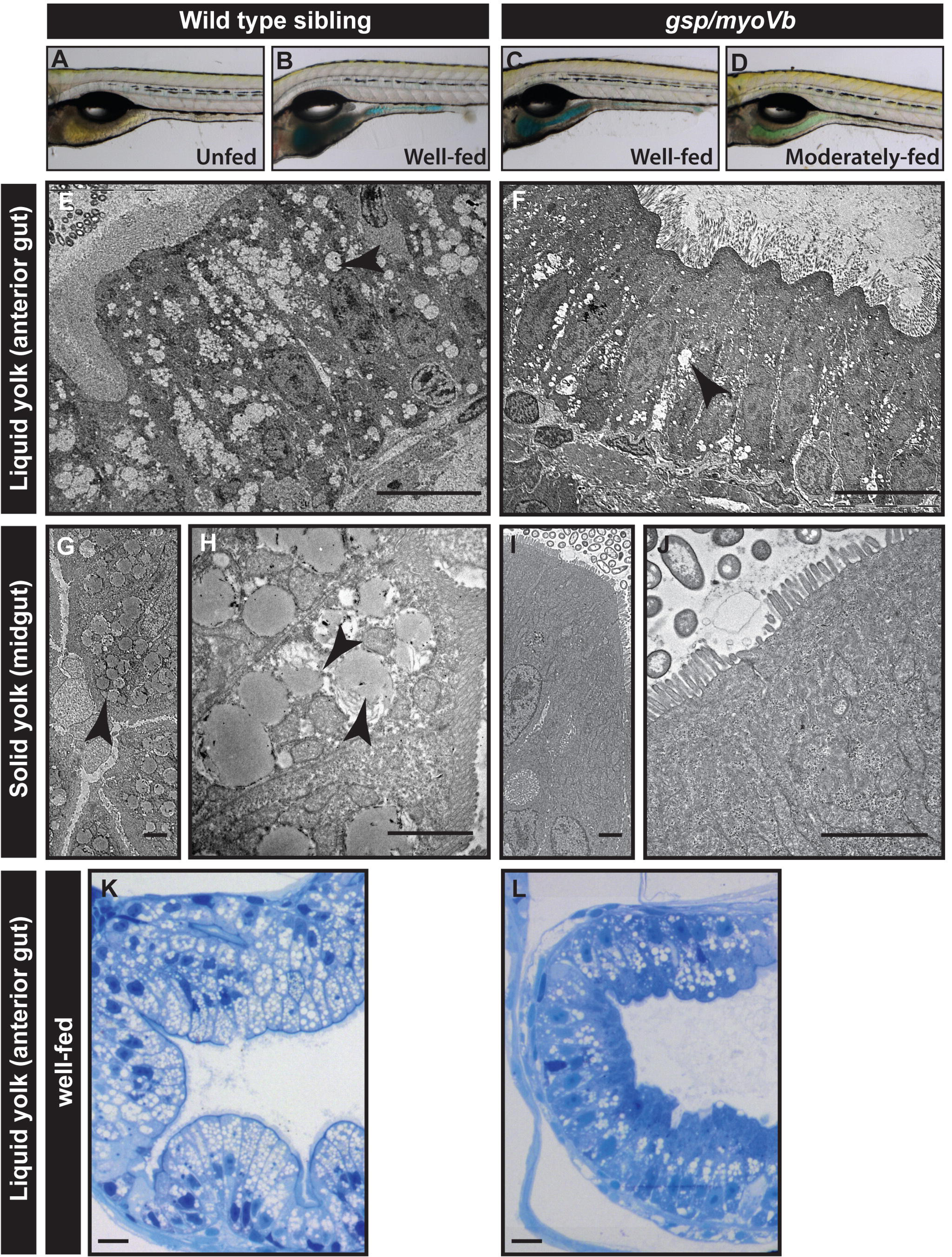
Lipid absorption is decreased in *gsp/myoVb* mutant larvae. Bright field images (A-D) of wild type (A,B) and mutant larvae (C,D), either unfed (A) or fed with coloured emulsified yolk (B-D). Electron micrograph of enterocytes from wild type siblings (E,G,H) and *gsp/myoVb* mutant larvae (F,I,J) fed with liquid yolk (E,F) and solid yolk (low magnification G,I; higher magnification H,J). Histological sections of the anterior gut of well-fed wild-type sibling (K) and *gsp/myoVb* (L) larvae. The micrographs of liquid yolk fed larval enterocytes were taken in the anterior part of the gut whereas the solid yolk fed larval enterocytes were imaged in the midgut. Note that while the absorption is low in the anterior gut in *gsp/myoVb* mutant larvae, it is almost absent in the midgut. The EM analysis was done on 3 wild type siblings and 3 mutants, out of 9 animals of each genotype, which were additionally analysed by histology (not shown). In the second set, 3 well-fed siblings and 4 well-fed mutants were preselected and used for histological analysis. The representative images for this set are presented in K and L. The black arrowheads point towards the lipid droplets in the enterocytes. Scale bars: in E, K and L corresponds to 10μm, in G, H, I, J correspond to 2μm

Our analysis convincingly proves that the loss of Myosin Vb function leads to a considerable reduction in lipid absorption.

## Discussion

The association of *myosin Vb* mutations with MVID has been clearly established over the last few years (Muller et al., 2008; Ruemmele et al., 2010). For quite some time, there was no animal model to study the pathophysiology of MVID or to understand the mechanistic basis of the cellular phenotype shown by enterocytes in MVID patients. Till date, depletion of Myosin Vb activity in Caco2 cells has been used as a strategy to recapitulate the cellular features of MVID (Ruemmele et al., 2010). Consequently, the exact function of *myosin Vb* in enterocytes has remained poorly understood. Very recently, while this paper was in revision, a *Myo5b* knockout mouse model recapitulating attributes of MVID has been published (Carton-Garcia et al., 2015). Here we report the zebrafish *gsp/myosin Vb* mutant as the first non-mammalian model for MVID.

Our analyses have revealed that the zebrafish *gsp/myoVb* mutant recapitulates morphological and cellular features of MVID. The gut morphology of *gsp/myoVb* mutant larvae exhibits drastic reduction in the intestinal folds. This is strikingly similar to the villus atrophy reported in human MVID patients. Furthermore, we show that in the absence of Myosin Vb function, enterocytes exhibit microvillus shortening and microvillus inclusions, pathognomonic features of MVID. Our data also show that the occurrence of microvillus inclusions is higher in the midgut than in the anterior gut. Some of the inclusions appear to be connected with the apical domain suggesting their apical origin. In addition to inclusion bodies, we observed the accumulation of secretory material in the apical half of the enterocytes in *gsp/myoVb* mutant larvae. These features are similar to those shown by human MVID patients. Our analyses thus indicate the conservation of the function of *myosin Vb* between fish and mammalian enterocytes. Although two other paralogues, *myosin Vaa* and *myosin Vc,* are expressed in the intestine, they do not seem to compensate for the loss of *myosin Vb* function.

In the mouse Myo5b knockout model, the reduction of blood glucose levels has been linked with watery diarrhoea and defects in absorption. We used the egg-yolk feeding assay (Walters et al., 2012) to directly investigate the effect of *gsp/myoVb* mutation on lipid absorption. While both wild type and mutant larvae ingested yolk, some of the mutant larvae showed reduced uptake of yolk as compared to the wild type. We observed considerable reduction in the lipid droplets in the mutant enterocytes. This reduction in lipid accumulation in mutant enterocytes might be a combined effect of decrease in absorption and, at least in some cases, due to less ingestion by the mutant larvae.

One of the novel findings of this study is the effect of loss of *myoVb* on the shape of enterocytes. Our analysis shows that wedge-shaped cells — apically broader and basally narrower — are associated with the intestinal folds in zebrafish larvae. In the absence of Myosin Vb function, the cells remain apically narrower and basally broader. We propose that the function of Myosin Vb is important to achieve the cell shape change, which is one of the driving forces underlying intestinal fold formation. The effect on cell shapes warranted analysis of enterocyte polarity. However, we did not observe any gross effect on the apico-basal polarity of the enterocytes. It appears that the effect on the cell polarity is manifested at a subtler level as mislocalisation of molecules like CD36, Na^+^/K^+^ ATPase and transferrin receptor have been reported earlier (Muller et al., 2008; Thoeni et al., 2014). While the gross apical-basal polarity seems fine at early larval stages, it is possible that the effect on polarity is secondary and becomes apparent at a later developmental stage. Further investigation is needed to resolve this issue.

We have previously reported that the decrease in epidermal cell size in the *gsp/myoVb* mutant is causally linked to cell proliferation (Sonal et al., 2014). Intriguingly, we did not observe any obvious changes in either cell number or proliferation in the mutant gut (JS, CP and MS; unpublished observations). Thus, the gut and epidermis seem to exhibit different responses to the same mutation indicating that these responses might be driven by architectural and/or physiological constraints on the tissue.

MVID is a fatal genetic disease. The therapies available include parenteral nutrition and bowel transplants (Oliva et al., 1994; Bunn et al., 2000; Ruemmele et al., 2004). Long-term parenteral nutrition may lead to complications such as bacterial infections, cholestatic liver disease and renal complications. In the absence of a suitable animal model, the progress on identifying small molecules that might help to alleviate the disease condition has not been possible so far. Zebrafish, due to its ex-utero development, high fecundity and transparent embryos, has emerged as a favourite organism in drug screening and discovery. Several zebrafish mutants and transgenic lines recapitulate human disease conditions. These strains have been proven important in screening chemical libraries to identify small molecules that would alleviate disease conditions. The zebrafish *gsp/myosin Vb* mutant reported here would be an invaluable tool for screening such libraries to identify compounds that will help manage the disease better and improve the lives of MVID patients.

## Experimental Procedures

### Fish strains

Three alleles of *goosepimples (gsp)* viz. *gsp^NS042^, gsp^K38B^* and *gsp^AT21^* were used for the morphological observations of the gut. However, all the analysis presented in this paper was done using *gsp^NS042^* allele. The transgenic line for *cldb:lyn-egfp* (Haas and Gilmour, 2006) was crossed to *gsp^NS042^* to obtain EGFP labelling of intestinal epithelium in *gsp/myoVb* mutants. For in situ hybridisation, embryos from the *albino* strain were used. For zebrafish maintenance and experimentation, the guidelines recommended by the Committee for the Purpose of Control and Supervision of Experiments on Animals (CPCSEA), Govt. of India, were followed.

### RT-PCR and in situ hybridisation

To check gene expression, intestines dissected out from 4 dpf larvae were used. RNA was extracted using a trizol (Invitogen, USA) based method. cDNA synthesis was carried out using up to 1 μg RNA using a cAMV reverse transcription kit (Invitrogen, USA). The following primers were used for RCR: *myosin-Vaa* (Forward: 5’gaacaaggagaaccgttcca3’; Reverse: 5’gtacgcaggagaaccaggag3’), *myosin-Vb* (Forward: 5’ acgagagacaacgatatcag; Reverse: 5’cttgttgagttgacgatttgg), *myosin-Vc* (Forward: 5’ acgggctcaagggttagaaa3 ‘; Reverse: 5’gcttgaagctgctcgttctc3’) and *β-actin* (Forward: 5’aaggccaacagggaaaagat3’; Reverse: 5’ aagtggtctcgtggataccg3’)

Using primers, a specific region of *myosin Vb* (NM_001161632.1; bp 2967-3866) was amplified and cloned into pCR TOPOII (Invitrogen, USA). Templates were linearised to synthesise probes using either T7 or SP6 RNA polymerase from DIG RNA Labelling Kit (Roche, Switzerland). In situ hybridisations were performed using a protocol described earlier with a few modifications (Schulte-Merker, 2002).

### Immunohistology

For whole intestine staining larvae were first fixed overnight at 4°C in 4% paraformaldehyde (PFA). After PBS washes the digestive tracts of the larvae were dissected out. The intestinal tissue was permeabilized with 0.8% PBT (0.8% Triton X-100 in phosphate buffer), blocked with 10% Normal Goat Serum (Jackson ImmunoResearch; USA) in PBT for 2 hours.

For staining intestinal sections GFP positive larvae were fixed in 4% PFA and kept overnight at 4°C. For E-cadherin staining larvae were fixed in Bouin’s fixative. The fixed larvae were upgraded to 30% sucrose and embedded in 14 cryomatrix blocks. 16 μ thin sections were cut using a cryotome (Leica, Germany). The sections were placed on poly-L-lysine coated slides. The slides were air-dried and the matrix was removed by distilled water and PBS washes. These sections were blocked with 10% Normal Goat Serum for 3 hours.

The whole intestinal tissue and cryosections were incubated overnight in anti-E-cadherin antibody (1:100; BD Transduction Labs, USA), anti-GFP antibody (1:200; Torrey Pines Biolabs, USA), anti-Lgl2 antibody (1:400; Sonawane et al., 2009), Alexa Fluor 594 conjugated WGA (5 µg/ml; Invitrogen, USA) or Rhodamine Phalloidin (1:40; Invitrogen, USA) in 1% NGS in PBT. After PBT washes, whereever appropriate, Alexa 488 (1:250; Invitrogen, USA) and Cy3 (1:750; Jackson ImmunoResearch, USA) conjugated secondary antibodies against mouse and rabbit were used in 1% NGS in PBT for 4 hours. The tissue or sections were washed in PBT, post-fixed in 4% PFA for 30 minutes and mounted in glycerol or vectashield (Vector Labs, USA) for imaging.

### Egg yolk feeding assay

This assay was performed as previously described (Walters et al., 2012) with a few modifications. 20 larvae each of the mutants and siblings were placed in 2 ml E3 buffer in separate wells in a 6 well plate. Either 600μl raw or 600 mg of boiled egg yolk was mixed well with E3 to obtain a homogenous emulsion of 2 ml egg yolk stock. Whenever essential 100μl food colour (GFC, batch no. 040/14, India), containing 1% brilliant blue FCF, propylene glycol, potable water and permitted diluent, was added to the stock. One ml of this stock was added to each of the two wells containing larvae to get 10% egg yolk solution in E3. The larvae were incubated with yolk for 3 hours, split into 2 groups of 10 larvae each, one of which was fixed for electron microscopy and the other for histology. The control group was left unfed and treated in the same way.

### Histology and electron microscopy

For histology, 6 dpf larvae were fixed in 4% PFA at room temperature for 30 minutes and then overnight at 4^0^C. They were dehydrated in ethanol and embedded in Epon-Araldite. 1μ sections were cut with a wet glass knife on a Leica Ultracut UC6 Ultramicrotome. Sections were lifted onto 1% gelatin coated slides and stained with Richardson’s Stain (Richardson et al., 1960) at 60^0^C for 2 minutes followed by thorough washing with water. Slides were mounted with DPX and imaged on a Zeiss Axioskop 2 plus, (Zeiss, Germany) with a Nikon Digital Sight DS-Fi2 (Nikon, Japan). The animals stained for whole mount in situ hybridisation were processed the same way as described above but sections were cut at 4μm thickness, mounted in glycerol and imaged using AxioCam MRc5 on Zeiss Apotome microscope.

For electron microscopy, 6 dpf larvae were fixed in PFA and glutaraldehyde 2.5% each in 0.1 M sodium cacodylate buffer (Electron Miroscopy Sciences Cat#15949) at room temperature for a half hour followed by overnight incubation at 4^0^C. They were washed 3 times in 0.1 M phosphate buffer (PB) at pH 6.2, post-fixed in cold 1% osmium tetroxide (Sigma 60H0150, USA) in 0.1 M PB at pH 6.2 for 45 minutes, washed thrice in distilled water, en bloc stained in 0.5% aqueous uranyl acetate (Electron Microscopy Sciences, USA) for 1 hour, again washed 3 times with distilled water, dehydrated in ethanol and embedded in Epon-Araldite. 70–100 nm sections were cut with a diamond knife (DiATOME, USA) on a Leica UC6 ultramicrotome (Leica, Germany) collected on slot grids, post-stained with lead citrate and imaged in a Zeiss Libra 120 TEM (Zeiss, Germany).

### Image acquisition and processing

For imaging immunostainings of whole intestines, the samples were mounted in 80% glycerol by placing a coverslip on them. Whole mounts and Vectashield (Vector Laboratories, USA) mounted cryosections of intestine were imaged using the Zeiss LSM 510 Meta with EC Plan-Neofluar 40X/1.30 oil objective (Zeiss, Germany). 1024 x1024 image dimensions were used, with an averaging of 4.

For imaging phenotypes, the embryos were treated with MESAB and mounted on Methyl Cellulose gel. Imaging of morphological phenotype and in situ hybridisation reaction was done on Zeiss SteREO Discovery with AxioCam (Zeiss, Germany).

Either ImageJ or ZEN Light Edition 2009 was used for image processing and analysis.

### Morphometric measurements of enterocytes, estimation of microvillus inclusions and data analysis

Confocal scans of cryosections of anterior, mid and posterior gut stained for lyn-EGFP were used for measurements of the central height, apical width and basal width of enterocytes. For each enterocyte a confocal slice was selected where the cell appeared the broadest. To quantify the cellular dimensions, lines corresponding to the height, apical and basal width were traced and measured using Measure tool of Image J. Around 90 cells from 7-9 larvae were analysed for wild type sibling and *gsp/myoVb* mutants each.

For estimation of the relative proportions of inclusion bodies, isolated intestines were phalloidin stained. Confocal stacks of whole mounts of the same frame size were taken, one each for the ventral wall of the anterior gut and the proximal midgut of the same animal (Zeiss 510 meta, 63× NA1.4, 1.5× zoom, 1024×1024 pixels, scaling X,Y: 0.093μm, scaling Z: 0.419 μm, 11-30 stacks). Actin rings showing a central clearing were counted from an equal number of slices of equal thickness from the two stacks of the same animal using the cell counter plugin on ImageJ. Repetitive counting was avoided by keeping track of inclusions in previous stacks. The data was collected from 5-6 animals from three different experiments.

For wild type siblings, the lengths of a total 137 microvilli were measured from 49 enterocytes from the midgut of 4 animals using ImageJ. For mutants, we estimated length from 95 microvilli from 33 midgut enterocytes of 3 animals.

The data was plotted using MS-Excel or by using http://boxplot.tyerslab.com/ and compared by a paired or unpaired Student’s t-test where appropriate using MS-Excel.

## Acknowledgements

We thank Dr. Kalidas Kohale, Sunny Gotarne and Sandip Shinde for maintenance of the fish facility. We acknowledge the help of Seema Shirolikar, Lalit Borde in electron microscopy and Tekchand Gaupale, Prajakta Dhage in cryosectioning.

## Competing Interests Statement

The authors declare no competing financial interests.

